# JaponicusDB: Rapid deployment of a model organism database for an emerging model species

**DOI:** 10.1101/2021.09.23.461587

**Authors:** Kim M. Rutherford, Midori A. Harris, Snezhana Oliferenko, Valerie Wood

## Abstract

The fission yeast *Schizosaccharomyces japonicus* has recently emerged as a powerful system for studying the evolution of essential cellular processes, drawing on similarities as well as key differences between *S. japonicus* and the related, well-established model *Schizosaccharomyces pombe*. We have deployed the open-source, modular code and tools originally developed for PomBase, the *S. pombe* model organism database (MOD), to create JaponicusDB (www.japonicusdb.org), a new MOD dedicated to *S. japonicus*. By providing a central resource with ready access to a growing body of experimental data, ontology-based curation, seamless browsing and querying, and the ability to integrate new data with existing knowledge, JaponicusDB supports fission yeast biologists to a far greater extent than any other source of *S. japonicus* data. JaponicusDB thus enables *S. japonicus* researchers to realise the full potential of studying a newly emerging model species, and illustrates the widely applicable power and utility of harnessing reusable PomBase code to build a comprehensive, community-maintainable repository of species-relevant knowledge.

## Introduction

In recent years the fission yeast *Schizosaccharomyces japonicus* has emerged as a powerful system for studying evolutionary cell biology, via comparison to the related, well-established model eukaryote *Schizosaccharomyces pombe*. Although these two fission yeasts carry out conserved cell-level processes using similar gene complements, substantial physiological differences in the mechanisms underpinning cell growth and division (Aoki et al., 2011; Yam et al., 2011; Gu et al., 2015; Makarova et al., 2016), physiology (Okamoto et al., 2013; Kinnaer et al., 2019), and metabolism (Bulder, 1963; Kaino et al., 2018; Makarova et al., 2020) provide invaluable opportunities to study how processes accomplished by universally conserved gene products may diverge. Indeed, *S. pombe* and *S. japonicus* may be regarded together as a composite model system in which conserved processes, gene products, and associated phenotypes can be compared side by side (Gu and Oliferenko, 2015; Russell et al., 2017; Oliferenko, 2018). Furthermore, *S. japonicus* can be used as a standalone model organism for studying biological processes not apparent or readily tractable in other model yeasts, such as nuclear envelope breakdown and reassembly (Yam et al., 2013; Pieper et al., 2020), cellular geometry scaling (Gu and Oliferenko, 2019) and quorum sensing (Gómez-Gil et al., 2019).

The current trajectory of *S. japonicus* research also aptly illustrates how the demands placed on species-specific data resources grow and change as an organism becomes established as a model. To date, the *S. japonicus* genome sequence and associated computationally generated annotation available from existing species-neutral data aggregators and repositories, such as the INSDC databases (https://www.insdc.org/), Ensembl Genomes (Howe et al., 2020), UniProtKB/TrEMBL (The UniProt Consortium, 2020), and FungiDB (Basenko et al., 2018), has sufficed for the early stages of research in this model organism. As a model system matures, however, researchers move beyond simple exploratory screens to designing more ambitious research programs that generate heterogeneous gene-specific molecular data for entire processes — a stage that *S. japonicus* research is now reaching. To accommodate growing bodies of literature and data, and to realise the full potential of coordinated studies investigating the divergent biology of fission yeasts, the accumulated results of *S. japonicus* experiments must be made readily available to the research community in the expertly curated and integrated state provided only by a model organism database (MOD; Oliver et al. 2016; Lipshitz 2021).

To address the urgent need for MOD infrastructure and services for *S. japonicus*, we have created JaponicusDB using the suite of open-source modular, customizable tools and code originally developed for PomBase, the *S. pombe* MOD (Lock et al. 2018 and Harris et al. in this issue). The PomBase database system was designed to facilitate re-use for emerging model species and comprises an online curation environment (Canto; Rutherford et al. 2014), a curation database using the Chado schema (Mungall et al., 2007), and code to import data and generate a website that features intuitive displays, a versatile query system, a genome browser (JBrowse; Buels et al. 2016), and support for daily data updates. Here, we describe the initial configuration and population of JaponicusDB and report on its current status. We summarise a round of manual curation in which we corrected and updated gene structure predictions, named genes, improved ortholog detection, and provided a greatly improved corpus of Gene Ontology (GO; The Gene Ontology Consortium 2000, 2021) annotation. Because community curation using Canto has proven successful in enhancing *S. pombe* literature curation (Lock et al., 2020), we have launched an analogous community curation approach for *S. japonicus* as a core part of JaponicusDB.

Now that both PomBase and JaponicusDB use the same intuitive database system and curation environment, all fission yeast researchers can easily carry out numerous activities that would not otherwise be feasible. Researchers can curate detailed data from small-scale experiments, including phenotypes, modifications, molecular functions, interactions, and processes; display data from publication-based curation rapidly and conveniently; query curated data via simple, intuitive tools; and compare lists of genes identified in specific experiments with comprehensively curated lists in an intuitive and meaningful way. The rapid deployment of JaponicusDB showcases how reusing the PomBase system brings immediate benefits to a model organism community in exchange for very reasonable input funds and effort.

## Methods

JaponicusDB shares a codebase with PomBase, which is used to build the database, load the data and run the website. All database-specific differences are captured in configuration files. The entire codebase is available from the PomBase GitHub organisation (https://github.com/pombase/), and the JaponicusDB configuration files and manual curation in a separate GitHub organisation (https://github.com/japonicusdb/). Now that the initial setup is complete, JaponicusDB can be maintained almost entirely by editing files stored in GitHub repositories; anyone with a GitHub account can thus be authorised to make corrections and other changes. The database and website are automatically updated daily using the latest code, ontology versions, configuration and data files. Problems detected during this process are recorded to publicly visible log files to allow remote users to investigate issues.

### Database initialisation

The first step in each daily update is to initialise a PostgreSQL database with an empty Chado schema (Mungall et al., 2007) and populate it with the required ontologies [Gene Ontology (GO; The Gene Ontology Consortium 2000, 2021), the Fission Yeast Phenotype Ontology (FYPO; Harris et al. 2013), the Sequence Ontology (SO; Eilbeck et al. 2005), the chemical ontology ChEBI (Hastings et al., 2016), the protein modification ontology PSI-MOD (Montecchi-Palazzi et al., 2008), and the Relations Ontology (RO; https://oborel.github.io/)].

### Preparing datasets

We prepared data files for the first JaponicusDB load as described below. As noted above, with the exception of external files from InterPro and GOA, all annotation files are now maintained in GitHub repositories.

### Genome

The *S. japonicus* genome contigs (EMBL:KE651166–KE651197; Rhind et al. 2011) and mitochondrial genome (EMBL:AF547983; Bullerwell et al. 2003) were downloaded from the European Nucleotide Archive (ENA; Harrison et al. 2021). Genome locus tags were repurposed to provide systematic identifiers, and identifiers (SJAGMIT_01–SJAGMIT_37) were minted for the mitochondrial genome.

### Transposons and LTRs

All transposons, transposon-related fragments and LTRs reported by Rhind et al. (2011) were added to the sequence contigs, assigned IDs in the range SJATN_00001–SJATN_00265, and loaded into the *S. japonicus* JBrowse instance.

### Orthologs

UniProtKB (The UniProt Consortium, 2020) records were downloaded using a query for taxon ID 402676 (the sequenced strain). Gene names applied by UniProtKB or via ENA submissions were imported. These were augmented by automated transfer of *S. pombe* gene names and products from PomBase where there is a one-to-one ortholog. Automatically transferred names are updated daily, with a configurable provision to override any unsuitable gene names or product descriptions. This configuration file also supplies manually curated names and product descriptions for paralogs, which we resolved based on synteny, and for genes with complicated orthology relationships, for which we used protein family membership to guide our decisions.

S. pombe *orthologs.* The *S. japonicus*–*S. pombe* orthologs identified by Rhind et al. (2011) for 4302 gene products were imported. These predictions were supplemented with orthologs from Ensembl Compara (Herrero et al., 2016), bringing the number of *S. japonicus* proteins with identified *S. pombe* orthologs to 4504. To find missing divergent orthologs, we identified proteins that are conserved between *S. pombe* and other species but did not have identified orthologs in *S. japonicus*, and devised gene-specific search strategies using a combination of FASTA (Pearson and Lipman, 1988), JackHMMER (Johnson et al., 2010), PSI-BLAST (Altschul et al., 1997), TBLASTN (Gertz et al., 2006), and synteny. This procedure also identified missing genes and gene structures that required revision.

#### Human and budding yeast orthologs

In PomBase, orthologs between *S. pombe* and *Saccharomyces cerevisiae* (budding yeast) and between *S. pombe* and human have been manually curated over twenty years using multiple ortholog prediction methods and tailored search strategies to provide complete and highly accurate coverage of known orthologs, including many not identified by automated methods (Wood, 2005; Hu et al., 2011). We inferred human and budding yeast orthologs by transferring orthologs manually curated by PomBase for *S. pombe* genes to the orthologous *S. japonicus* genes. These were supplemented with orthologs from Compara for proteins that have no *S. pombe* ortholog but are conserved in other species.

### Families and domains

Protein domain information from InterPro version 85 (Blum et al., 2021), together with coiled-coil and low-complexity regions from Pfam version 34.0 (El-Gebali et al., 2019), were imported using existing PomBase Chado loading code. Transmembrane domains were predicted using TMHMM (Krogh et al., 2001).

### Gene Ontology

GO annotations were obtained by downloading the GOA UniProt file produced by the Gene Ontology Annotation (GOA) project at EBI (Huntley et al., 2015), and then filtering for NCBI taxon ID 4897 or 402676. The loading script checks the GOA UniProt file (located at ftp://ftp.ebi.ac.uk/pub/databases/GO/goa/UNIPROT/goa_uniprot_all.gaf.gz) for updates upon each run. For the first JaponicusDB load, we used the GOA file from July 2021, which contains 36878 *S. japonicus* annotations.

The filtering protocol used by PomBase to retain only relevant and non-redundant automated annotation, in which GO annotations are filtered at two stages, was applied (Lock et al., 2018). Prior to loading into Chado, the GOA annotations were filtered to exclude any derived from false positive or otherwise inapplicable InterPro2GO (Finn et al. 2017, https://github.com/geneontology/go-site/blob/master/metadata/gorefs/goref-0000002.md) and keyword (https://github.com/geneontology/go-site/blob/master/metadata/gorefs/goref-0000041.md, https://github.com/geneontology/go-site/blob/master/metadata/gorefs/goref-0000043.md) mappings, or phylogeny-based transfer (Gaudet et al., 2011), and any GO terms that are flagged by GO as not usable for direct annotation, or as inapplicable due to taxon restrictions (Deegan née Clark et al., 2010; The Gene Ontology Consortium, 2021); after filtering, 24159 annotations were loaded.

To enhance the *S. japonicus* GO annotation set, *S. pombe* GO annotations were transferred to *S. japonicus* orthologs, and manual annotations from literature curated in Canto were added. The annotations loaded into Chado were then subjected to the second round of filtering, which removes redundant non-experimental GO annotation.

The JaponicusDB GO pipeline also supports configurable filtering that can be used in cases where paralogs have diverged (e.g. one or both of a paralogous pair has undergone neofunctionalisation or sub-functionalisation), making the *S. pombe* annotation inappropriate for *S. japonicus*; with automated annotation transfer suppressed, the paralogs can be manually annotated. Annotation from PomBase is also not transferred for any gene–term combination that contradicts a manually curated negated (NOT) annotation in JaponicusDB (e.g. Mid1 GO:1902408). Finally, for known proteins conserved in other species but absent from *S. pombe*, GO annotations were made manually by inference from experimentally characterised orthologs. As of September 2021, JaponicusDB has 29589 GO annotations.

Identification of missing genes and revision of gene structures *Protein-coding genes.* During manual ortholog assignment, we observed that a number of genes at syntenic locations did not display any clear sequence similarity. Inspecting the structures of these genes revealed errors, which were corrected. All gene structure editing was done using the Artemis genome annotation tool (Rutherford et al., 2000). Gene structures annotated in the published assembly without methionines were either trimmed to the appropriate methionine, or revised to include a missing N-terminal exon. Gene structure errors reported in publications were also corrected (Makarova et al., 2016, 2020).

We created a list of *S. pombe* genes that are broadly conserved in single copy throughout eukaryotes (predominantly one-to-one to human) using the PomBase advanced search and imported the result list into JaponicusDB using the identifier mapping feature (Harris et al. in this issue). This query combination provided a list of conserved proteins present in *S. pombe* but absent from *S. japonicus*. Since we expected most of these genes to be present in *S. japonicus*, we used TBLASTN to perform directed searches against the *S. japonicus* genome, using the *S. pombe* proteins as input, coupled with manual inspection and gene structure curation in Artemis to identify highly spliced genes at syntenic locations.

#### Non-coding RNAs

Annotated features imported with the genome sequence included 347 tRNA genes and 17 rRNA genes, but no other non-coding RNAs. We used tRNAscan-SE (Chan and Lowe, 2019) to confirm the tRNA complement, identifying three additional tRNA genes (SJAG 08001–SJAG 08003), and as a source of codon information for the product descriptions. Other missing non-coding RNAs were imported from RNAcentral (The RNAcentral Consortium, 2019).

### Website code and documentation

The JaponicusDB website uses the PomBase website code (Lock et al., 2018), configured for *S. japonicus*. In brief, the Chado database is processed nightly to build a Docker container which holds a complete instance of the website, including all documentation, HTML, code, and data.

All documentation for JaponicusDB is maintained in Markdown format in a GitHub repository and the source files are shared with PomBase where possible.

### Canto setup and literature import

The Canto community curation tool was designed to support different species and datatypes via simple configuration adjustments (Rutherford et al., 2014). As deployed for *S. japonicus*, Canto currently supports annotating phenotypes using FYPO, alleles and genotypes, protein modifications, genetic and physical interactions, qualitative gene expression, and GO, including associated annotation extensions (Huntley et al., 2014; Lock et al., 2018) and metadata. A PubMed search using a set of *S. japonicus*-related keywords retrieved 179 publications, which were imported into Canto along with all citation details needed to populate Canto and the website. Publications containing gene-specific data were flagged using Canto’s literature triage function, which allows users to classify papers by type (e.g., curatable, review, methods), and then assign them to authors for full curation. The triage procedure also flagged 91 false positives, most of which were publications describing a “japonicus” species in a genus other than *Schizosaccharomyces*. We have refined the PubMed search keywords based on these results, so future Canto publication updates will include fewer false positives.

## Results

### Adding value to imported data

#### Adding and revising gene structures

The original gene models for *S. japonicus* were built using aligned RNA-seq transcripts from *S. pombe*, *S. cryophilus* and *S. octosporus* orthologous groups compared to protein sequence (Rhind et al., 2011). Gene models with discrepancies were then manually reviewed to provide a set of high-quality gene predictions. However, continual revisions are required to make corrections and keep gene structures in line with published knowledge.

In this work, we used previously published information (Makarova et al., 2016) and detailed orthology inspection to revise 36 gene structures, illustrated in Figure 1. We identified 18 additional conserved genes (SJAG 07000–SJAG 07017) by searching for small, universally conserved proteins absent from the predicted gene set. In eight cases, we replaced erroneous gene predictions with genes correctly identified in an alternative reading frame. Newly identified genes, include Atp19, an F-type ATPase subunit for which no gene structure previously existed (Figure 1A), and the recombination protein Rec7, for which the originally predicted structure was completely replaced by a new structure (Figure 1B). For SJAG 03830, the predicted structure was revised but not completely replaced (Figure 1C), allowing the gene product to be identified as the ubiquinol-cytochrome-c reductase complex subunit Qcr10, previously thought absent from *S. japonicus*.

**Fig. 1.**
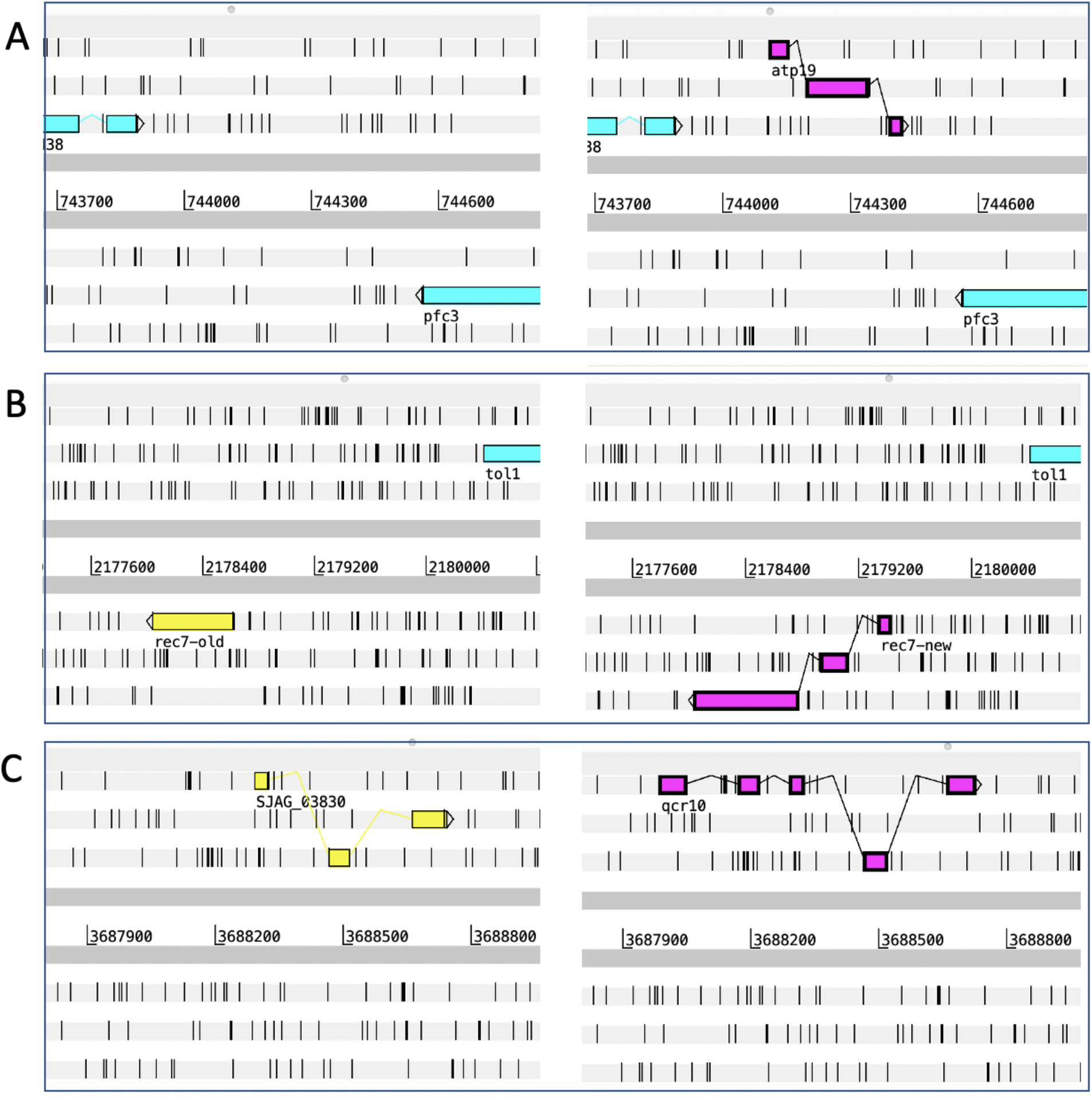
Examples of updated *S. japonicus* gene structures, illustrated using screenshots from the Artemis genome annotation tool (Rutherford et al., 2000). For each gene, the original genomic annotation is shown on the left and the revised annotation on the right. A) *atp19*, a newly identified gene; B) *rec7*, for which a new structure completely replaces the original structure; C) *qcr10*, for which the original structure was revised but not completely replaced.

Using tRNAscan-SE (Chan and Lowe, 2019), we added three genes to the tRNA complement, and added product descriptions specifying codons to the 325 nuclear-encoded tRNAs. Additional non-coding RNAs including snRNAs, snoRNAs, and telomerase RNA were imported from RNAcentral (The RNAcentral Consortium, 2019).

#### Identifying distant orthologs

Accurate detection of orthologs, i.e. genes in different species related by vertical descent (Fitch, 1970), between two proteomes allows for comprehensive comparative study of the biological processes in two species. Orthologs provide a framework for reconstructing the evolutionary events that have given rise to observed biological differences. Thorough ortholog inventories require the targeted detection of distant orthologs and the identification of the orthologous relationship type (i.e., one-to-one, one-to-many, or many-to-many). Ortholog inventories also facilitate the accurate identification of lineage specific gene losses, species-specific genes, and protein family expansions.

In this study, we identified 67 previously undetected distant orthologs between *S. pombe* and *S. japonicus*, 25 of which are also conserved in human (Supplementary Table S1). In two cases ortholog detection depended on the revision of existing gene structures and in 18 cases, finding new genes. Candidates for a further 17 orthologs (including the kinetochore regulator Meikin, the kinetochore protein Mis19, and the spindle component Dms1) are recorded as high confidence based on a one-to-one relationship between *S. pombe* and other species and syntenic location between *S. pombe* and *S. japonicus*, although sequence similarity has not yet been detected. These regions are currently being reviewed for possible gene prediction errors. Finally, some additional ortholog calls from the automated pipelines were revised to include in-paralogs and remove out-paralogs.

#### Gene names and product descriptions

##### Gene names

In consultation with *S. japonicus* researchers, we have assigned standard gene names in JaponicusDB, using unified nomenclature to make conserved loci readily recognisable and to avert potential naming conflicts with other species. Only 101 gene names were imported from UniProtKB for *S. japonicus*. Names for 3471 protein-coding genes were assigned by importing *S. pombe* names from PomBase for all clear one-to-one orthologs. Manual curation of 119 exact or near-exact duplicate proteins (ribosomal proteins, histones, translation elongation factors) enabled us to assign names for syntenic orthologs. Finally, 184 gene names were assigned manually to *S. japonicus* specific families, one-to-many, and many-to-one paralogs. Overall, JaponicusDB now provides standard names for 3875 protein-coding genes (of 4896 total). The remit of the fission yeast Gene Naming Committee (GNC; https://www.pombase.org/submit-data/gene-names) has expanded to cover *S. japonicus* as well as *S. pombe* gene names; the GNC will approve all new gene names in both species.

##### Gene product descriptions

Informative gene product descriptions allow users to browse proteomes effectively, scan input and output lists from experiments, and review genome contents. Gene product descriptions for 4251 protein-coding genes were automatically transferred from one-to-one orthologs curated by PomBase. A further 645 descriptions were manually curated based on one-to-many, many-to-one, or many-to-many orthologs or protein family descriptions. JaponicusDB now provides gene product labels that have been reviewed by fission yeast experts, and accurately reflect current knowledge for 4896 protein-coding genes.

Our analysis and curation pipeline has improved gene product descriptions dramatically compared to previously available data. UniProtKB provides descriptions for protein gene products, but over 99% of *S. japonicus* proteins remain in the unreviewed TrEMBL database and therefore have only automated descriptions. The quality of TrEMBL’s inferred descriptions is generally high, but coverage is incomplete. Notably, we have provided informative product labels for 522 proteins described as “uncharacterized protein” in UniProtKB. Many of these are broadly conserved, well-studied proteins identified and annotated via our distant ortholog detection or manual curation pipelines. For example, we found five proteins involved in TOR signalling and three cytochrome oxidase assembly factors.

The JaponicusDB pipeline updates imported gene names and product descriptions upon each run. Simple configuration files maintained in GitHub repositories specify manually curated names and descriptions that take precedence over automated imports.

#### Extending GO annotation coverage and specificity

To generate a Gene Ontology (GO; The Gene Ontology Consortium 2000, 2021) annotation dataset for *S. japonicus*, we imported computationally generated annotations from the GOA UniProt (Huntley et al., 2015), and added new annotations from the PomBase ortholog pipeline, manual inferences for genes without *S. pombe* orthologs, and experimental data from literature manually curated in Canto. We applied the filtering protocol used by PomBase (Lock et al., 2018) to retain only relevant and non-redundant automated annotation. Annotations manually curated from the literature in Canto take precedence over any automatically transferred annotations (for example, as noted in Methods, manual negated annotations suppress import of contradictory automated annotations). The JaponicusDB GO annotation procedures thus support robust transfer of large annotation sets, while also allowing the fine tuning of specific annotation to capture known biological differences (reviewed in Gu and Oliferenko 2015; Russell et al. 2017; Oliferenko 2018; also see references cited in Introduction). The resulting GO annotation dataset comprises 29589 annotations as of September 2021.

PomBase maintains subsets of GO, known as “GO slims”, of selected biologically meaningful terms from each branch of GO — Molecular Function (MF), Biological Process (BP), and Cellular Component (CC) — that are used to summarise the functional capabilities of *S. pombe* (Lock et al., 2018; Wood et al., 2019). JaponicusDB uses the PomBase GO slims for the same purposes, illustrating the value added by our GO curation procedures. GO slim analysis shows that annotation breadth has significantly increased compared to the GO data originally available from GOA despite an overall decrease in annotation number (from 36878 to 29512). For each aspect of GO, more genes are annotated to terms in the GO slim (MF coverage increased from 3087 to 3652, BP from 3238 to 4187, and CC from 2999 to 4578 genes), and the number of genes that either have no annotation, or are annotated but not covered by the GO slim, is correspondingly decreased (Figure 2). Notably, we have increased annotation coverage in several areas of active *S. japonicus* research. For example, the annotation increases for “membrane organization” and “lipid metabolism” in BP and “endoplasmic reticulum” (ER) and “mitochondrion” in CC are highly relevant to comparative studies of respiration and membrane and lipid biology. In MF, there is a large increase in annotations to “hydrolase activity”, including 21 gene products annotated to “lipid metabolism”. Newly annotated examples in this set include the mitochondrial cardiolipin-specific phospholipase Cld1, the acyl-coenzyme A thioesterase The4, and SJAG 00199, a mitochondrial DDHD family phospholipase. The relatively new GO term “molecular adaptor activity”, relevant to research in organelle organisation, also has increased annotations including many newly described protein- and organelle-to-membrane adaptors.

**Fig. 2.**
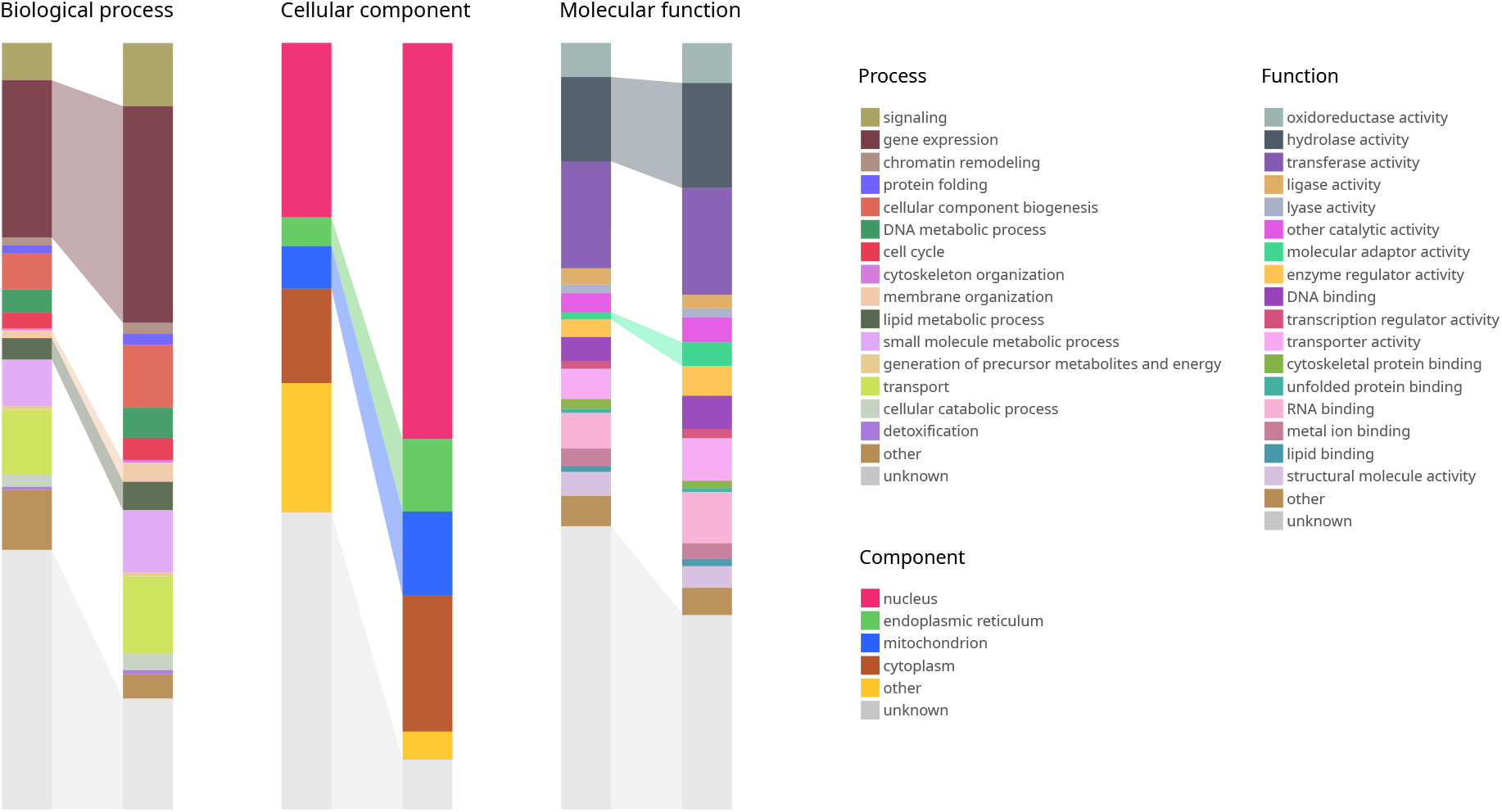
Effect of curation on *S. japonicus* GO annotation coverage. Comparison of the distribution of annotations for all *S. japonicus* protein-coding genes (4896 total) to selected high-level terms in each aspect of GO between the UniProt GOA annotation file (“before”; left-hand columns) and the JaponicusDB GO annotation set (“after”; right-hand columns). The number of annotated proteins has increased for most Molecular Function (MF), Biological Process (BP), and Cellular Component (CC), with associated decreases in the proportions of proteins annotated as “unknown”. Highlighted blocks represent GO terms relevant to processes that are intensively studied in *S. japonicus*: “gene expression”, “membrane organization” and “lipid metabolism” in BP; “endoplasmic reticulum” and “mitochondrion” in CC; and “hydrolase activity” and “molecular adaptor activity” in MF. Images were generated using QuiLT (Harris et al., in this issue), in which only one term per gene can be included for display. If a gene is annotated to more than one GO term, one is selected according to a set order of precedence. Here, terms are arranged by order of precedence in the charts and key for each GO aspect.

Annotation depth, defined as increased distance from the root node in an ontology graph, provides a measure of the biological specificity of the annotations. Our work has increased annotation depth in at least one GO aspect (Molecular Function, Biological Process or Cellular Component) for 4312 proteins (>88% of the total), and in all 3 aspects for 1385 (>28%) (Figure 3).

**Fig. 3.**
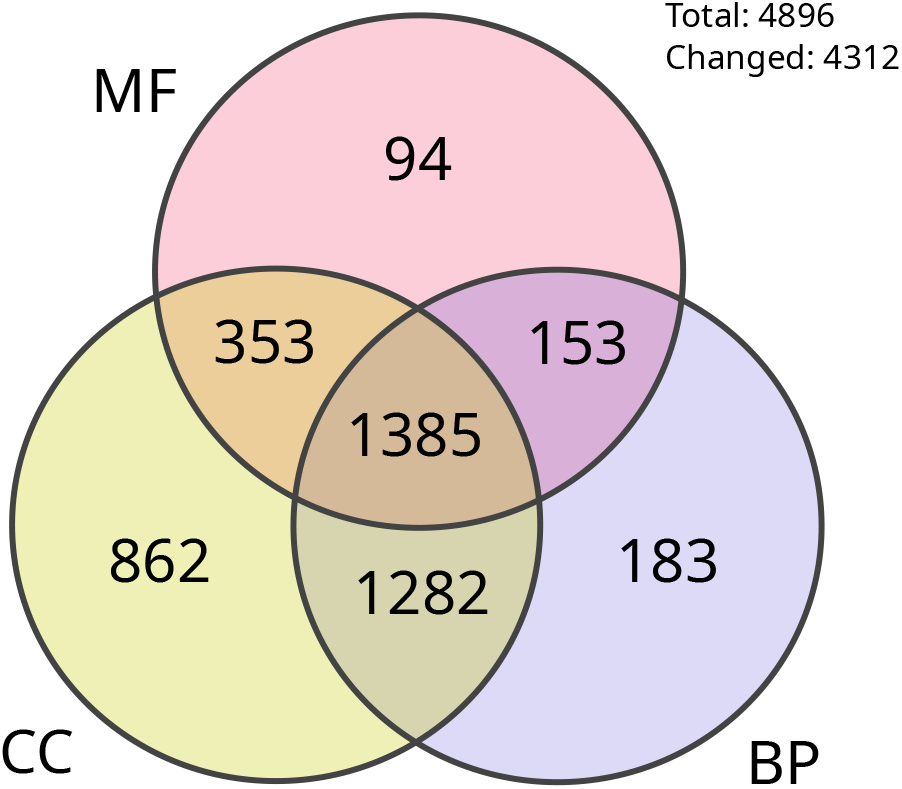
*S. japonicus* proteins with increased GO annotation specificity. GO annotation specificity, defined as distance from the ontology root, was measured for each aspect of GO and for each of the 4896 proteins in *S. japonicus*. The Venn diagram shows the number of proteins that have more specific annotations in the JaponicusDB GO annotation set than in the UniProt GOA annotation file for one or more GO aspect(s) (MF, Molecular Function; BP, Biological Process; CC, Cellular Component).

#### Manual literature curation in Canto

Using Canto’s literature triage function, we have manually classified the 179 publications found in our initial PubMed query (see Methods). After discarding spurious matches, we identified 31 papers containing gene-specific data suitable for curation, as well as 39 papers in other categories. The latter reflect the status of *S. japonicus* as an emerging model, and include publications describing wild type features and cell composition, methods and reagents, or phylogenetic studies, as well as reviews. The 31 curatable papers were assigned to the authors (or to a Canto administrator) for curation, and 23 are now fully curated in JaponicusDB (see https://www.japonicusdb.org/reference_list/community). To date, manual curation has supplied 135 experimentally supported GO annotations and 168 phenotype annotations.

### Website

The JaponicusDB website, modelled on PomBase, provides convenient access to all molecular and cell biological data curated for *S. japonicus*. Like PomBase, JaponicusDB includes pages for each gene, publication, and genotype annotated, as well as for ontology terms. JaponicusDB also uses the same simple, advanced, and peptide motif search tools as PomBase. To facilitate comparison between *S. japonicus* and *S. pombe*, the “Ortholog” section of each gene page includes reciprocal links between PomBase and JaponicusDB. On both sites, orthologs can also be retrieved via the advanced search or via an ID mapping tool. The JaponicusDB front page (Figure 4) presents news, database usage hints, publications recently curated by the community, and “Research Spotlight” panels, which feature graphical abstracts from recent papers (one of the most popular features of PomBase). An instance of the genome browser JBrowse (Buels et al., 2016) currently displays the *S. japonicus* genome and annotated features, and will host any available high-throughput sequence-based datasets. Online documentation describes all page contents, search capabilities, and how to contribute to JaponicusDB (also see Lock et al. 2018 and Harris et al. in this issue). JaponicusDB uses Google Analytics to monitor website usage.

**Fig. 4.**
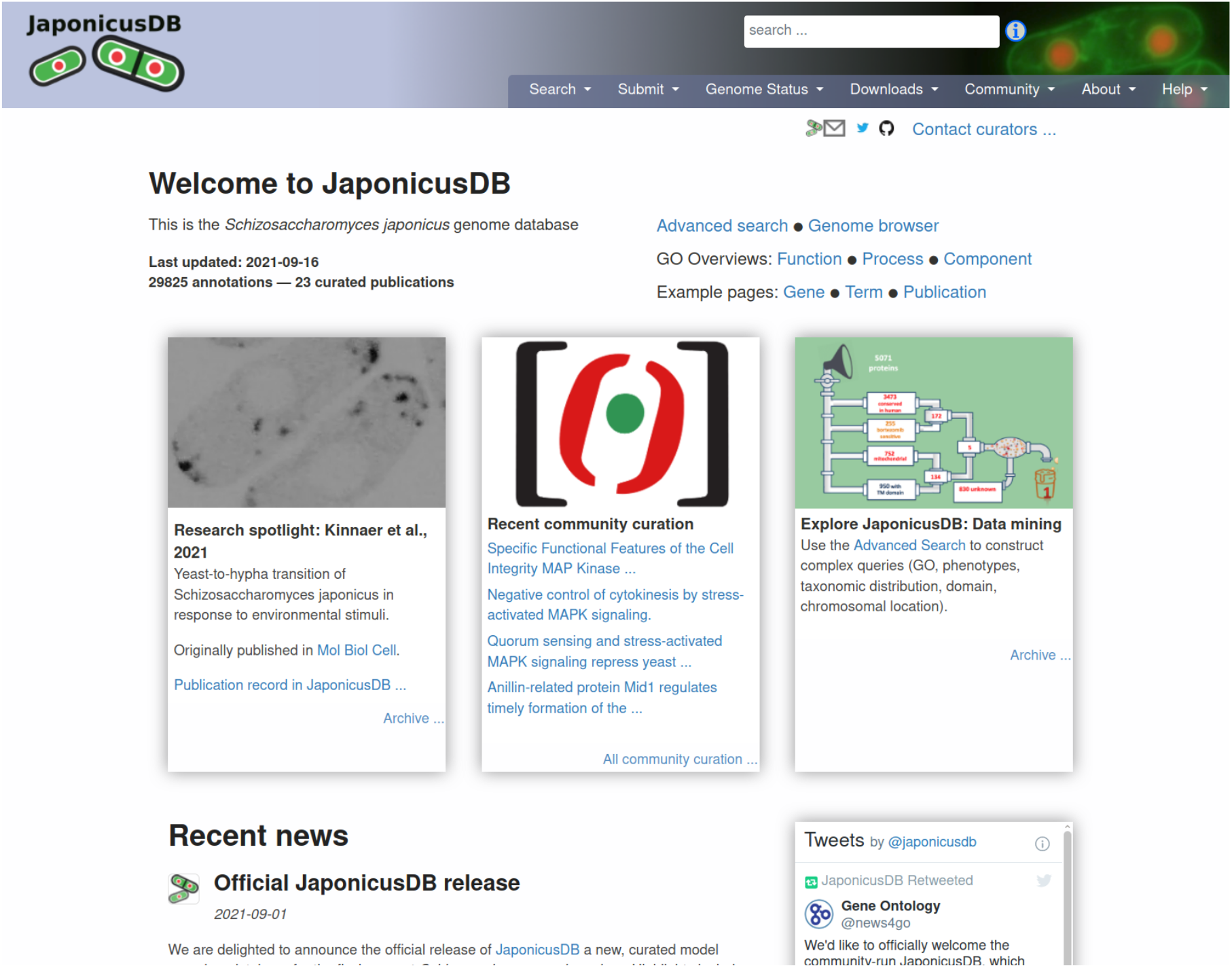
The JaponicusDB front page.

## Conclusions

For several decades, fission yeast research has been a mainstay of progress in cell and molecular biology, due to pioneering work on cell cycle regulation, as well as recent innovative experimentation in additional areas such as chromosome segregation, epigenetics and cytokinesis, using *S. pombe*. The growing amount and sophistication of research on *S. pombe*’s “sister” species *S. japonicus* provides a prime opportunity to add evolutionary biology to the roster of topics that can be fruitfully investigated using fission yeast.

We have launched JaponicusDB to enable *S. japonicus* researchers — and the wider scientific community — to realise the full promise of experiments in an emerging model system. JaponicusDB gathers and interconnects diverse types of information from many sources to create a single resource that provides coherent, intuitive access to rich, interconnected data sets. As a fully functional MOD, JaponicusDB supports robust genome-wide data interrogation and mining at any level of detail, experimental planning, hypothesis generation, and interpretation of results, thereby meeting research needs not addressed by any other system.

### PomBase code re-use

We have re-used the modular, configurable open-source code underlying PomBase, enabling us to deploy a curation database, a Canto instance, and a functioning, intuitive website for JaponicusDB with modest developer and curator input. (We estimate that essential development took the equivalent of approximately 1 FTE for six weeks, and we devoted a similar amount of time to curation, i.e. 12 FTE-weeks total.) For updates, the PomBase system relies primarily on data and configuration files stored (with version control) in GitHub repositories, making future maintenance straightforward. Because JaponicusDB uses PomBase code, both databases will simultaneously deploy any new features developed by the PomBase team.

### Community literature curation

The Canto community literature curation system enables bench biologists to contribute directly to the knowledge integrated into JaponicusDB. Although Canto is also being used for phenotype curation in FlyBase (Larkin et al., 2021) and pathogen–host interaction phenotypes in PHI-Base (Urban et al., 2020), JaponicusDB has embraced Canto for community curation at a uniquely early stage in its development. The curatable literature corpus is small at present, but over two thirds of suitable papers have already been curated. *S. japonicus* researchers can use community curation to meet data dissemination objectives, helping ensure timely FAIR (Findable, Accessible, Interoperable and Reusable; Wilkinson et al. 2016) sharing of new information. To enhance engagement with JaponicusDB and interactions among community members, a mailing list has been set up (https://mailman.kcl.ac.uk/mailman/listinfo/japonicus-list).

### Community MOD management

To ensure that JaponicusDB meets its users’ needs in the future, we will invite experts in relevant areas, including *S. japonicus* biology and database curation and maintenance, to form a Scientific Advisory Board (SAB). The SAB will collaborate with PomBase staff to mobilise volunteers from the *S. japonicus* community to carry out routine updates and maintenance, and will guide future fission yeast MOD developments that arise from *S. japonicus* research. PomBase staff will provide training and advice to community volunteers in configuring the website and JBrowse track metadata, Canto administration, editing gene structures, and other maintenance tasks. This group will also arrange to disseminate data regularly from JaponicusDB to external resources, e.g. GO annotations to the GO Consortium repository.

### Curation to support *S. japonicus* research

Our review of the *S. japonicus* genome highlights the value that manual curation adds to sequence feature annotation, ortholog identification, and functional annotation, and therefore to all subsequent usage of these data in experimental and predictive studies. As a result, JaponicusDB can provide more comprehensive, experimentally informed complements of several types of genome-scale data than were previously available via data aggregators. We note that finding genes previously thought to be absent from *S. japonicus* has significant consequences for accurately representing molecular and cellular processes within the species as well as comparative studies of how these processes have evolved.

Gene structures for most genomes are derived by automated prediction pipelines, often incorporating information from homology and synteny. Although multiple methods are frequently used, many errors will persist even for intensively curated model species in which all gene predictions were manually inspected prior to publication. For example, since publication of the *S. pombe* genome, PomBase has revised >200 gene structures; similarly, WormBase has updated several thousand for C. elegans (pers. comm. Paul Davis, WormBase). Every genome-scale analysis depends upon comprehensive, manually refined gene structures (and often other sequence features) for accuracy, reproducibility, and relevance.

Our work also illustrates how biologists may benefit directly from manual ortholog curation. For example, we have identified *S. japonicus* orthologs of cytochrome c oxidase subunits Cox7 and Cox9, ATP synthase subunit Atp19, mitochondrial alpha-ketoglutarate dehydrogenase Ymr1, and ubiquinol-cytochrome-c reductase complex subunit Qcr10, all of which are important for fission yeast researchers studying the differences in central carbon metabolism between *S. pombe* and *S. japonicus*.

Future updates to ortholog predictions, gene structures, names, and product descriptions, are bound to reveal more examples where comprehensive, curated knowledge is indispensable for understanding biological processes, avoiding misinterpretation, and making reliable cross-species comparisons

### Rapid MOD deployment to support emerging model species

The rapid establishment of JaponicusDB promises to have beneficial repercussions well beyond the fission yeast community. First, simply having a full *S. japonicus* MOD available to facilitate and support research makes reliable information available to underpin comparative studies and data integration, not only between *S. japonicus* and *S. pombe*, but throughout all eukaryotes.

Perhaps of even greater consequence, JaponicusDB represents a proof of concept, demonstrating that a small group of people can easily deploy PomBase code to produce a comparable system for any other species. We and others have noted (Oliver et al., 2016; Alliance of Genome Resources Consortium, 2019; Lipshitz, 2021) that complete genome sequences continue to accumulate, but annotation even of sequence features, let alone any other associated data, cannot keep pace. A growing number of increasingly diverse research communities require creative approaches to data infrastructure to take advantage of the exploratory opportunities that a genome sequence offers, and to integrate genome-scale molecular data and methods with small-scale results from new or extant published literature. Our experience with JaponicusDB exemplifies one way forward: the PomBase system can be redeployed by a small, motivated community to create a resource that can be maintained within the scope of time and skills available to researchers, using funding allocated to data dissemination.

## Data Availability

As described in detail above, JaponicusDB uses the open-source PomBase code base available from the PomBase GitHub organisation (https://github.com/pombase). JaponicusDB configuration and data files are available from the JaponicusDB GitHub organisation (https://github.com/japonicusdb).

JaponicusDB data can be viewed directly at https://www.japonicusdb.org, query results provide data download options, and data can be downloaded in bulk from the website (see https://github.com/japonicusdb.org/datasets and https://www.https://github.com/japonicusdb.org/data).

## Acknowledgements

We are grateful to the Wellcome Trust for allowing S. O. to redirect funds from a Senior Investigator Award (see below) to establish JaponicusDB. We thank Nick Rhind and Li-Lin Du for advice and discussions about the *S. japonicus* genome sequence, Blake Sweeney of the RNAcentral Consortium for help with non-coding RNA gene annotation, and the *S. japonicus* researchers who have contributed community curation (Sophie Martin, Makoto Kawamukai, Elisa Gomez-Gil, and Ying Gu).

## Funding

This research was funded in whole, or in part, by the Wellcome Trust [Grant number 103741/Z/14/Z to S. O.]. For the purpose of open access, the author has applied a CC BY public copyright licence to any Author Accepted Manuscript version arising from this submission.

S. O. also acknowledges funding from Wellcome Trust (Investigator Award in Science [220790/Z/20/Z]) and BBSRC [BB/T000481/1]. The Francis Crick Institute receives its core funding from Cancer Research UK (FC001002), the UK Medical Research Council (FC001002), and the Wellcome Trust (FC001002). PomBase is supported by the Wellcome Trust [218236/Z/19/Z to Juan Mata].

## Conflicts of interest

None declared.

## References

Alliance of Genome Resources Consortium. The Alliance of Genome Resources: Building a modern data ecosystem for model organism databases. Genetics, 213(4):1189–1196, 2019.

S. F. Altschul, T. L. Madden, A. A. Schäffer, J. Zhang, Z. Zhang, W. Miller, and D. J. Lipman. Gapped BLAST and PSI-BLAST: a new generation of protein database search programs. Nucleic Acids Res, 25(17):3389–3402, 1997.

K. Aoki, H. Hayashi, K. Furuya, M. Sato, T. Takagi, M. Osumi, A. Kimura, and H. Niki. Breakage of the nuclear envelope by an extending mitotic nucleus occurs during anaphase in Schizosaccharomyces japonicus. Genes Cells, 16(9):911–926, 2011.

E. Y. Basenko, J. A. Pulman, A. Shanmugasundram, O. S. Harb, K. Crouch, D. Starns, S. Warrenfeltz, C. Aurrecoechea, C. J. Stoeckert, J. C. Kissinger, D. S. Roos, and C. Hertz-Fowler. FungiDB: An integrated bioinformatic resource for fungi and oomycetes. J Fungi (Basel), 4(1), 2018.

M. Blum, H.-Y. Chang, S. Chuguransky, T. Grego, S. Kandasaamy, A. Mitchell, G. Nuka, T. Paysan-Lafosse, M. Qureshi, S. Raj, L. Richardson, G. A. Salazar, L. Williams, P. Bork, A. Bridge, J. Gough, D. H. Haft, I. Letunic, A. Marchler-Bauer, H. Mi, D. A. Natale, M. Necci, C. A. Orengo, A. P. Pandurangan, C. Rivoire, C. J. A. Sigrist, Sillitoe, N. Thanki, P. D. Thomas, S. C. E. Tosatto, C. H. Wu, A. Bateman, and R. D. Finn. The InterPro protein families and domains database: 20 years on. Nucleic Acids Res, 49(D1):D344–D354, 2021.

R. Buels, E. Yao, C. M. Diesh, R. D. Hayes, M. Munoz-Torres, G. Helt, D. M. Goodstein, C. G. Elsik, S. E. Lewis, L. Stein, and I. H. Holmes. JBrowse: a dynamic web platform for genome visualization and analysis. Genome Biol, 17:66, 2016.

C. J. E. A. Bulder. On respiratory deficiency in yeasts. PhD thesis, Technological University, Delft, Delft, The Netherlands, 1963.

C. E. Bullerwell, J. Leigh, L. Forget, and B. F. Lang. A comparison of three fission yeast mitochondrial genomes. Nucleic Acids Res, 31(2):759–768, 2003.

P. P. Chan and T. M. Lowe. tRNAscan-SE: Searching for tRNA genes in genomic sequences. Methods Mol Biol, 1962:1–14, 2019.

J. I. Deegan née Clark, E. C. Dimmer, and C. J. Mungall. Formalization of taxon-based constraints to detect inconsistencies in annotation and ontology development. BMC Bioinformatics, 11:530, 2010.

K. Eilbeck, S. E. Lewis, C. J. Mungall, M. Yandell, L. Stein, R. Durbin, and M. Ashburner. The Sequence Ontology: a tool for the unification of genome annotations. Genome Biol, 6 (5):R44, 2005.

S. El-Gebali, J. Mistry, A. Bateman, S. R. Eddy, A. Luciani, S. C. Potter, M. Qureshi, L. J. Richardson, G. A. Salazar, A. Smart, E. L. L. Sonnhammer, L. Hirsh, L. Paladin, D. Piovesan, S. C. E. Tosatto, and R. D. Finn. The Pfam protein families database in 2019. Nucleic Acids Res, 47 (D1):D427–D432, 2019.

R. D. Finn, T. K. Attwood, P. C. Babbitt, A. Bateman, P. Bork, A. J. Bridge, H.-Y. Chang, Z. Dosztányi, S. El-Gebali, M. Fraser, J. Gough, D. Haft, G. L. Holliday, H. Huang, X. Huang, I. Letunic, R. Lopez, S. Lu, A. Marchler-Bauer, H. Mi, J. Mistry, D. A. Natale, M. Necci, G. Nuka, C. A. Orengo, Y. Park, S. Pesseat, D. Piovesan, S. C. Potter, N. D. Rawlings, N. Redaschi, L. Richardson, C. Rivoire, A. Sangrador-Vegas, C. Sigrist, I. Sillitoe, B. Smithers, S. Squizzato, G. Sutton, N. Thanki, P. D. Thomas, S. C. E. Tosatto, C. H. Wu, I. Xenarios, L.-S. Yeh, S.-Y. Young, and A. L. Mitchell. InterPro in 2017-beyond protein family and domain annotations. Nucleic Acids Res, 45(D1):D190–D199, 2017.

W. M. Fitch. Distinguishing homologous from analogous proteins. Syst Zool, 19(2):99–9113, 1970.

P. Gaudet, M. S. Livstone, S. E. Lewis, and P. D. Thomas. Phylogenetic-based propagation of functional annotations within the Gene Ontology consortium. Brief Bioinform, 12 (5):449–462, 2011.

E. M. Gertz, Y.-K. Yu, R. Agarwala, A. A. Schäffer, and S. F. Altschul. Composition-based statistics and translated nucleotide searches: improving the TBLASTN module of BLAST. BMC Biol, 4:41, 2006.

E. Gómez-Gil, A. Franco, M. Madrid, B. Vázquez-Marín, M. Gacto, J. Fernández-Breis, J. Vicente-Soler, T. Soto, and J. Cansado. Quorum sensing and stress-activated mapk signaling repress yeast to hypha transition in the fission yeast Schizosaccharomyces japonicus. PLoS Genet, 15(5): e1008192, 2019.

Y. Gu and S. Oliferenko. Comparative biology of cell division in the fission yeast clade. Curr Opin Microbiol, 28:18–25, 2015.

Y. Gu and S. Oliferenko. Cellular geometry scaling ensures robust division site positioning. Nat Commun, 10(1):268, 2019.

Y. Gu, C. Yam, and S. Oliferenko. Rewiring of cellular division site selection in evolution of fission yeasts. Curr Biol, 25(9): 1187–1194, 2015.

M. A. Harris, A. Lock, J. Bähler, S. G. Oliver, and V. Wood. FYPO: the fission yeast phenotype ontology. Bioinformatics, 29(13):1671–1678, 2013.

P. W. Harrison, A. Ahamed, R. Aslam, B. T. F. Alako, J. Burgin, N. Buso, M. Courtot, J. Fan, D. Gupta, M. Haseeb, S. Holt, T. Ibrahim, E. Ivanov, S. Jayathilaka, V. B. Kadhirvelu, M. Kumar, R. Lopez, S. Kay, R. Leinonen, X. Liu, C. O’Cathail, A. Pakseresht, Y. Park, S. Pesant, N. Rahman, J. Rajan, A. Sokolov, S. Vijayaraja, Z. Waheed, A. Zyoud, T. Burdett, and G. Cochrane. The European Nucleotide Archive in 2020. Nucleic Acids Res, 49(D1): D82–D85, 2021.

J. Hastings, G. Owen, A. Dekker, M. Ennis, N. Kale, V. Muthukrishnan, S. Turner, N. Swainston, P. Mendes, and C. Steinbeck. ChEBI in 2016: Improved services and an expanding collection of metabolites. Nucleic Acids Res, 44 (D1):D1214–1219, 2016.

J. Herrero, M. Muffato, K. Beal, S. Fitzgerald, L. Gordon, M. Pignatelli, A. J. Vilella, S. M. J. Searle, R. Amode, S. Brent, W. Spooner, E. Kulesha, A. Yates, and P. Flicek. Ensembl comparative genomics resources. Database (Oxford), 2016, 2016.

K. L. Howe, B. Contreras-Moreira, N. D. Silva, G. Maslen, W. Akanni, J. Allen, J. Alvarez-Jarreta, M. Barba, D. M. Bolser, L. Cambell, M. Carbajo, M. Chakiachvili, M. Christensen, C. Cummins, A. Cuzick, P. Davis, S. Fexova, A. Gall, N. George, L. Gil, P. Gupta, K. E. Hammond-Kosack, E. Haskell, S. E. Hunt, P. Jaiswal, S. H. Janacek, P. J. Kersey, N. Langridge, U. Maheswari, T. Maurel, M. D. McDowall, B. Moore, M. Muffato, G. Naamati, S. Naithani, A. Olson, I. Papatheodorou, M. Patricio, M. Paulini, H. Pedro, E. Perry, J. Preece, M. Rosello, M. Russell, V. Sitnik, D. M. Staines, J. Stein, M. K. Tello-Ruiz, S. J. Trevanion, M. Urban, S. Wei, D. Ware, G. Williams, A. D. Yates, and P. Flicek. Ensembl Genomes 2020-enabling non-vertebrate genomic research. Nucleic Acids Res, 48(D1): D689–D695, 2020.

Y. Hu, I. Flockhart, A. Vinayagam, C. Bergwitz, B. Berger, N. Perrimon, and S. E. Mohr. An integrative approach to ortholog prediction for disease-focused and other functional studies. BMC Bioinformatics, 12:357, 2011.

R. P. Huntley, M. A. Harris, Y. Alam-Faruque, J. A. Blake, S. Carbon, H. Dietze, E. C. Dimmer, R. E. Foulger, D. P. Hill, V. K. Khodiyar, A. Lock, J. Lomax, R. C. Lovering, P. Mutowo-Meullenet, T. Sawford, K. V. Auken, V. Wood, and C. J. Mungall. A method for increasing expressivity of Gene Ontology annotations using a compositional approach. BMC Bioinformatics, 15:155, 2014.

R. P. Huntley, T. Sawford, P. Mutowo-Meullenet, A. Shypitsyna, C. Bonilla, M. J. Martin, and C. O’Donovan. The GOA database: Gene Ontology annotation updates for 2015. Nucleic Acids Res, 43(Database issue):D1057–1063, 2015.

L. S. Johnson, S. R. Eddy, and E. Portugaly. Hidden Markov model speed heuristic and iterative HMM search procedure. BMC Bioinformatics, 11:431, 2010.

T. Kaino, K. Tonoko, S. Mochizuki, Y. Takashima, and M. Kawamukai. Schizosaccharomyces japonicus has low levels of CoQ10 synthesis, respiration deficiency, and efficient ethanol production. Biosci Biotechnol Biochem, 82(6): 1031–1042, 2018.

C. Kinnaer, O. Dudin, and S. G. Martin. Yeast-to-hypha transition of Schizosaccharomyces japonicus in response to environmental stimuli. Mol Biol Cell, 30(8):975–991, 2019.

A. Krogh, B. Larsson, G. von Heijne, and E. L. Sonnhammer. Predicting transmembrane protein topology with a hidden Markov model: application to complete genomes. J Mol Biol, 305(3):567–580, 2001.

A. Larkin, S. J. Marygold, G. Antonazzo, H. Attrill, G. dos Santos, P. V. Garapati, J. L. Goodman, L. S. Gramates, G. Millburn, V. B. Strelets, C. J. Tabone, J. Thurmond, and the FlyBase Consortium. FlyBase: updates to the Drosophila melanogaster knowledge base. Nucleic Acids Res, 49(Database issue):D899–D907, 2021.

H. D. Lipshitz. The descent of databases. Genetics, 217(3), 2021.

A. Lock, K. Rutherford, M. A. Harris, J. Hayles, S. G. Oliver, J. Bähler, and V. Wood. PomBase 2018: user-driven reimplementation of the fission yeast database provides rapid and intuitive access to diverse, interconnected information. Nucleic Acids Research, 47(D1):D821–D827, 2018. doi: 10.1093/nar/gky961. URL http://dx.doi.org/10.1093/nar/gky961.

A. Lock, M. A. Harris, K. Rutherford, J. Hayles, and V. Wood. Community curation in PomBase: enabling fission yeast experts to provide detailed, standardized, sharable annotation from research publications. Database (Oxford), 2020:baaa028, 2020. ISSN 1758-0463. doi: 10.1093/database/baaa028. URL https://doi.org/10.1093/database/baaa028.

M. Makarova, Y. Gu, J.-S. Chen, J. R. Beckley, K. L. Gould, and S. Oliferenko. Temporal regulation of lipin activity diverged to account for differences in mitotic programs. Curr Biol, 26(2):237–243, 2016.

M. Makarova, M. Peter, G. Balogh, A. Glatz, J. I. MacRae, N. L. Mora, P. Booth, E. Makeyev, L. Vigh, and S. Oliferenko. Delineating the rules for structural adaptation of membrane-associated proteins to evolutionary changes in membrane lipidome. Curr Biol, 30(3):367–380.e8, 2020.

L. Montecchi-Palazzi, R. Beavis, P.-A. Binz, R. J. Chalkley, J. Cottrell, D. Creasy, J. Shofstahl, S. L. Seymour, and J. S. Garavelli. The PSI-MOD community standard for representation of protein modification data. Nat Biotechnol, 26(8):864–866, 2008.

C. J. Mungall, D. B. Emmert, and FlyBase Consortium. A Chado case study: an ontology-based modular schema for representing genome-associated biological information. Bioinformatics, 23(13):i337–346, 2007.

S. Okamoto, K. Furuya, S. Nozaki, K. Aoki, and H. Niki. Synchronous activation of cell division by light or temperature stimuli in the dimorphic yeast Schizosaccharomyces japonicus. Eukaryot Cell, 12(9):1235–1243, 2013.

S. Oliferenko. Understanding eukaryotic chromosome segregation from a comparative biology perspective. J Cell Sci, 131(14), 2018.

S. G. Oliver, A. Lock, M. A. Harris, P. Nurse, and V. Wood. Model organism databases: essential resources that need the support of both funders and users. BMC Biol, 14:49, 2016.

W. R. Pearson and D. J. Lipman. Improved tools for biological sequence comparison. Proc Natl Acad Sci U S A, 85(8): 2444–2448, 1988.

G. H. Pieper, S. Sprenger, D. Teis, and S. Oliferenko. ESCRT-III/Vps4 controls heterochromatin-nuclear envelope attachments. Dev Cell, 53(1):27–41.e6, 2020.

N. Rhind, Z. Chen, M. Yassour, D. A. Thompson, B. J. Haas, N. Habib, I. Wapinski, S. Roy, M. F. Lin, D. I. Heiman, S. K. Young, K. Furuya, Y. Guo, A. Pidoux, H. M. Chen, B. Robbertse, J. M. Goldberg, K. Aoki, E. H. Bayne, A. M. Berlin, C. A. Desjardins, E. Dobbs, L. Dukaj, L. Fan, M. G. FitzGerald, C. French, S. Gujja, K. Hansen, D. Keifenheim, J. Z. Levin, R. A. Mosher, C. A. Müller, J. Pfiffner, M. Priest, C. Russ, A. Smialowska, P. Swoboda, S. M. Sykes, M. Vaughn, S. Vengrova, R. Yoder, Q. Zeng, R. Allshire, D. Baulcombe, B. W. Birren, W. Brown, K. Ekwall, M. Kellis, J. Leatherwood, H. Levin, H. Margalit, R. Martienssen, C. A. Nieduszynski, J. W. Spatafora, N. Friedman, J. Z. Dalgaard, P. Baumann, H. Niki, A. Regev, and C. Nusbaum. Comparative functional genomics of the fission yeasts. Science, 332(6032):930–936, 2011.

J. J. Russell, J. A. Theriot, P. Sood, W. F. Marshall, L. F. Landweber, L. Fritz-Laylin, J. K. Polka, S. Oliferenko, T. Gerbich, A. Gladfelter, J. Umen, M. Bezanilla, M. A. Lancaster, S. He, M. C. Gibson, B. Goldstein, E. M. Tanaka, C.-K. Hu, and A. Brunet. Non-model model organisms. BMC Biol, 15(1):55, 2017.

K. Rutherford, J. Parkhill, J. Crook, T. Horsnell, P. Rice, M. A. Rajandream, and B. Barrell. Artemis: sequence visualization and annotation. Bioinformatics, 16(10):944–945, 2000.

K. M. Rutherford, M. A. Harris, A. Lock, S. G. Oliver, and V. Wood. Canto: an online tool for community literature curation. Bioinformatics, 30(12):1791–1792, 2014.

The Gene Ontology Consortium. Gene Ontology: tool for the unification of biology. Nat Genet, 25(1):25–29, 2000.

The Gene Ontology Consortium. The Gene Ontology resource: enriching a GOld mine. Nucleic Acids Res, 49(D1):D325–D334, 2021.

The RNAcentral Consortium. RNAcentral: a hub of information for non-coding RNA sequences. Nucleic Acids Res, 47(D1):D221–D229, 2019.

The UniProt Consortium. UniProt: the universal protein knowledgebase in 2021. Nucleic Acids Research, 49(D1): D480–D489, 11 2020. ISSN 0305-1048. doi: 10.1093/nar/gkaa1100. URL https://doi.org/10.1093/nar/gkaa1100.

M. Urban, A. Cuzick, J. Seager, V. Wood, K. Rutherford, S. Y. Venkatesh, N. D. Silva, M. C. Martinez, H. Pedro, A. D. Yates, K. Hassani-Pak, and K. E. Hammond-Kosack. PHI-base: the pathogen-host interactions database. Nucleic Acids Res, 48(D1):D613–D620, 2020.

M. D. Wilkinson, M. Dumontier, I. J. Aalbersberg, G. Appleton, M. Axton, A. Baak, N. Blomberg, J.-W. Boiten, L. B. da Silva Santos, P. E. Bourne, J. Bouwman, A. J. Brookes, M. Clark, T. Crosas, O. Dillo, I. Dumon, S. Edmunds, C. R. Evelo, T. Finkers, A. Gonzalez-Beltran, A. J. G. Gray, P. Groth, C. Goble, J. S. Grethe, J. Heringa, P. A. C. ’t Hoen, T. Hooft, R. Kuhn, R. Kok, J. Kok, S. J. Lusher, M. E. Martone, A. Mons, A. L. Packer, B. Persson, M. Rocca-Serra, P. Roos, R. van Schaik, S.-A. Sansone, E. Schultes, T. Sengstag, T. Slater, G. Strawn, M. A. Swertz, M. Thompson, J. van der Lei, E. van Mulligen, J. Velterop, A. Waagmeester, P. Wittenburg, K. Wolstencroft, J. Zhao, and B. Mons. The FAIR Guiding Principles for scientific data management and stewardship. Sci Data, 3:160018, 2016.

V. Wood. Schizosaccharomyces pombe comparative genomics; from sequence to systems, pages 233–285. Springer, Berlin, Heidelberg, 2005. ISBN 978-3-540-31480-6.

V. Wood, A. Lock, M. A. Harris, K. Rutherford, J. Bähler, and S. G. Oliver. Hidden in plain sight: what remains to be discovered in the eukaryotic proteome? Open Biol, 9(2): 180241, 2019.

C. Yam, Y. He, D. Zhang, K.-H. Chiam, and S. Oliferenko. Divergent strategies for controlling the nuclear membrane satisfy geometric constraints during nuclear division. Curr Biol, 21(15):1314–1319, 2011.

C. Yam, Y. Gu, and S. Oliferenko. Partitioning and remodeling of the Schizosaccharomyces japonicus mitotic nucleus require chromosome tethers. Curr Biol, 23(22):2303–2310, 2013.

